# A Cost-Effective CRISPR/Cas9-Based Method for Sequencing the Functional Antibody Kappa Chain cDNA from Hybridomas Expressing Aberrant Kappa Chain mRNAs

**DOI:** 10.1101/2023.06.08.544271

**Authors:** Armin Ahnoud, Chong He, Michael Strainic, Wenwu Zhai

## Abstract

One prerequisite for generating recombinant antibodies for therapeutic and non-therapeutic purposes from hybridoma clones is the reliable, efficient, and cost-effective sequencing of the hybridoma clone that produces the desired antibody. However, many hybridoma fusion partners produce aberrant endogenous mRNA transcripts most of which resemble kappa chains. These aberrant kappa chain mRNAs can interfere with or even prevent the determination of the functional murine antibody kappa chain cDNA sequences during the PCR amplification step. In this paper, we report the development of a rapid and cost-effective CRISPR/Cas9 based method to eliminate the aberrant endogenous sequence. We have demonstrated the effectiveness of this method by significantly reducing the number of an endogenous aberrant kappa chain transcript known as the aberrant SP2/0 kappa chain. This transcript is produced by the commonly used fusion partner known as SP2/0 which is a myeloma-derived cell line. Here, we first cloned and sequenced two hybridoma clones using this method (clones A1E7 and A9E11). Our results showed a 24 to 25 percent reduction of the aberrant chain sequences (method 1) from the sequencing results. We then optimized this method (method 2) and used it to clone and sequence a third hybridoma clone (clone 262). This optimized method allowed us to achieve an 88% reduction of the aberrant kappa chain sequences.

## 1.0 Introduction

The discovery of hybridoma technology by César Milstein and Georges J. F. Köhler in 1975 [1,2] revolutionized our ability to generate antibodies against almost any molecule. Today, hybridoma clones are usually generated by immunizing an animal (most commonly mice), isolating their B cells, and fusing them to a fusion partner. Furthermore, modern molecular cloning techniques have allowed the generation of chimeric and even humanized antibodies in a matter of weeks. One of the requirements to generate chimeric or humanized antibodies is the successful sequencing of the hybridoma clones. However, depending on the fusion partner, the hybridoma might also express high levels of non-functional endogenous transcripts [3]. These endogenous transcripts can interfere with obtaining the sequence of the functional transcript thus making the hybridoma sequencing process expensive, tedious, and laborious. This problem is mostly seen when sequencing the kappa chains. Most cell lines that are used as murine fusion partners are derived from the original p3-X63-Ag8.653 myeloma cell line [4]. This cell line expresses an endogenous and aberrant kappa chain mRNA that is not translated into a functional kappa chain protein because of a four nucleotide deletion at the VJ joining site which produces a stop codon after the complementary determining region (CDR) 3 [4]. Over the years, various strategies have been developed to sequence the antibody light and heavy chains. Methods that rely on next generation sequencing (NGS) have proven to be more successful [5]. However, such methods are usually expensive and require access to specialized instruments [6]. Traditionally, the genetic information of antibody heavy and light chains has been amplified from hybridoma cells via polymerase chain reaction (PCR) using degenerate primers [7]. However, these primer sets sometimes resulted in low success rates because of non-specific annealing or no annealing. In 2019, Meyer et al. reported the development of a new method by designing universal primers that could amplify all possible variable region sequences with a high success rate. This method is relatively simple, cost effective, and does not require specialized instruments [7].

The Clustered Regularly Interspaced Short Palindromic Repeats (CRISPR)/Cas9 system evolved as a bacterial defense mechanism against viruses and plasmids [5]. The system is composed of two main components. The first component is the RNA which itself is made of two types of RNAs: the CRISPR RNA (crRNA) which binds specifically to the double-stranded DNA (dsDNA); and the Trans-activating CRISPR RNA (TracrRNA) which binds to the crRNA. These two RNA transcripts can be combined into a single guide RNA (sgRNA). The second component is the Cas9 protein which binds to the TracrRNA or the TracrRNA component of the sgRNA. The binding of Cas9 to sgRNA results in a conformational change and therefore activates the Cas9 protein. Activated Cas9 then recognizes the protospacer adjacent motif (PAM) sequence. For *S. pyogenes* Cas9, the PAM sequence is a three nucleotide sequence (NGG) located on the target DNA immediately downstream of the sgRNA binding site. In the presence of a PAM sequence, Cas9 introduces a double-stranded break in the DNA. It has been previously demonstrated that CRISPR/Cas9 can cleave dsDNA in vitro [12]. Lastly, CRISPR/Cas9 has been used to cleave unwanted sequences in vitro for various applications. For example, for next generation RNAseq library preparations [13,14,15]; and for contamination-free PCR based viral RNA detection tests for diagnostic purposes [16].

In the past three decades, various strategies have been utilized to eliminate the aberrant endogenous kappa chain transcripts in hybridoma clones to obtain the functional kappa chain sequence. First, Juste et al. utilized restriction endonucleases to cleave the aberrant light chain rather than the functional one. These enzymes were applied after PCR amplification of the antibody variable genes [8]. Second, Yuan et al. utilized a panel of kappa primers in the presence of molar excess of a primer complementary to the CDR3 of the known aberrant chain sequence [9]. Third, Ostermeier et al. developed an antisense-directed RNase H against the aberrant light chain mRNA from hybridoma clones [10]. Lastly, Duan et al. developed a ribozyme specific for the aberrant VL gene and packaged it into a retroviral expression vector system in order to eliminate the endogenous aberrant light chain mRNA [11]. These strategies, although relatively effective, are laborious, time-consuming, and can sometimes be unreliable. To our knowledge, the CRISPR/Cas9 system has not been applied to hybridoma cloning and sequencing processes.

In this paper, we have developed a method to specifically cleave the SP2/0 endogenous kappa transcript using the CRISPR/Cas9 system via targeting its CDR3 region. This strategy also can be applied to any other endogenous transcript by modifying the sgRNA sequence. In method1, we CRISPR/Cas9 treated the cDNA after the RT-PCR; next, we performed the PCR reaction. In method2, we CRISPR/Cas9 treated the PCR amplicon, gel purified the uncleaved band (560bp), PCR amplified the purified product again and CRISPR/Cas9 digested it again (double cut). Next, we gel purified, cloned, and sequenced the uncleaved band. The summary of both methods is shown in figure1.

**Figure 1:**
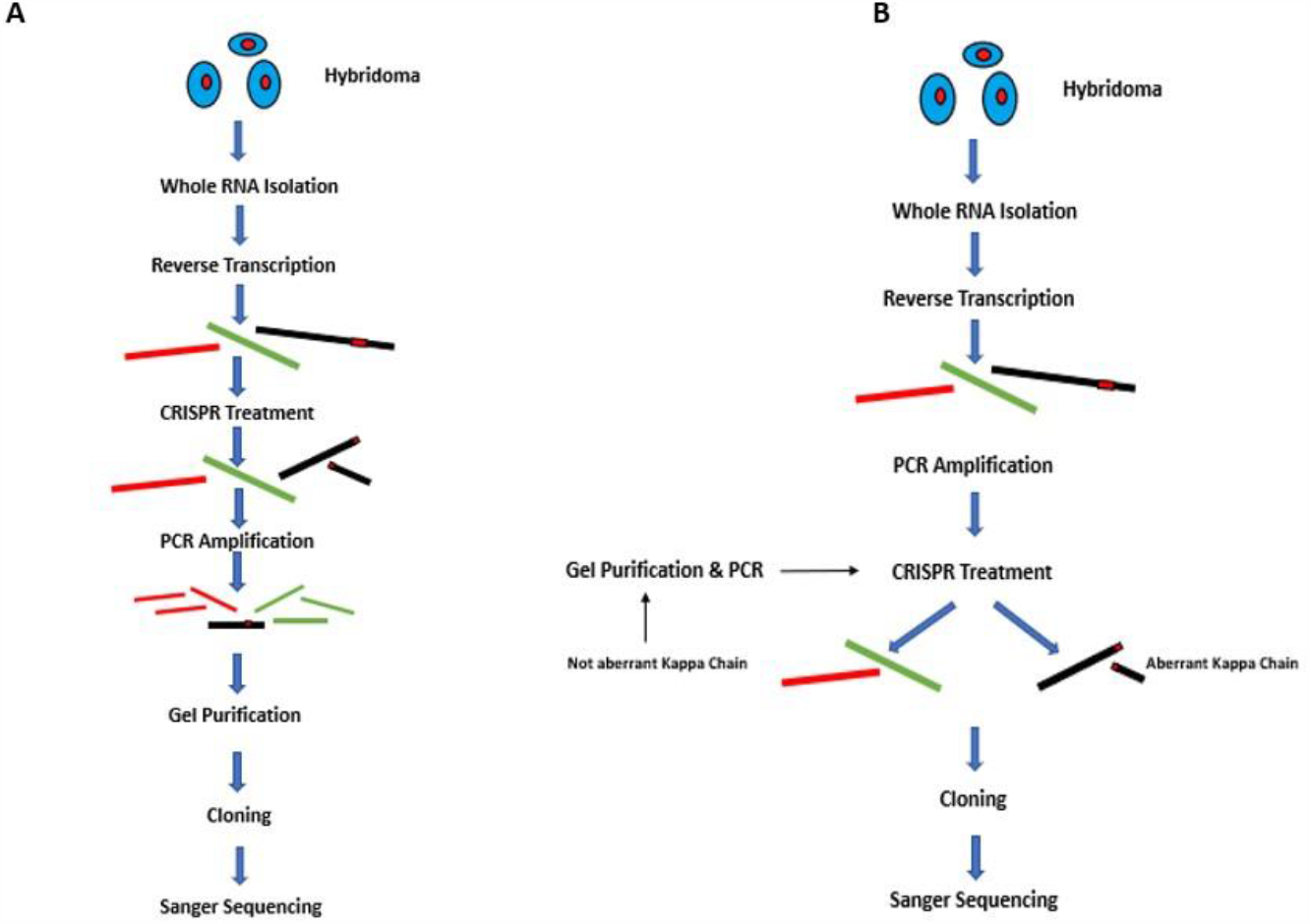
A is a schematic of method 1. B is a schematic of the optimized method (method2).

## 2.0 MATERIALS AND METHODS

### 2.1 Hybridoma Generation

The hybridoma clones were generated in-house by fusing mouse (Balb/c) B cells to SP2/0 cell line. The spleens and lymph nodes were first harvested and the B cells were isolated using mouse B cell isolation kit (StemCell, 19854A). Next, the cells were fused to the SP2/0 cells using BTX ECM 2001 + electrofusion machine (BTX, 45-2048) using ECM 2001+ electrofusion protocol (protocol 1016) [17]. Hypoxanthine-Aminopterin-Thymidine (HAT) selection was performed using semi-solid medium D (StemCell, 03804) and the colonies were picked after 10-14 days using ClonePix2 (Molecular Devices). The isotypes of antibodies generated from hybridoma clones were determined using Pierce mouse Ig isotyping kit (Thermo Fisher 26178). The selected hybridomas produced IgG.

### 2.2 sgRNA Design

In order to minimize the off-target effect of the sgRNA, we used the CDR3 of the SP2/0 aberrant chain as the target for sgRNA binding. Custom Alt-R® CRISPR-Cas9 guide RNA (IDT, https://www.idtdna.com/site/order/designtool/index/CRISPR_SEQUENCE) was used to design and order the sgRNA. (5’-ACCTATTACTGTCAGCACATTAGGGAGCTTACACGTTCGGAGGGGGGACCAAGCTGGAAATAAAACG - 3’) was used as the input sequence.

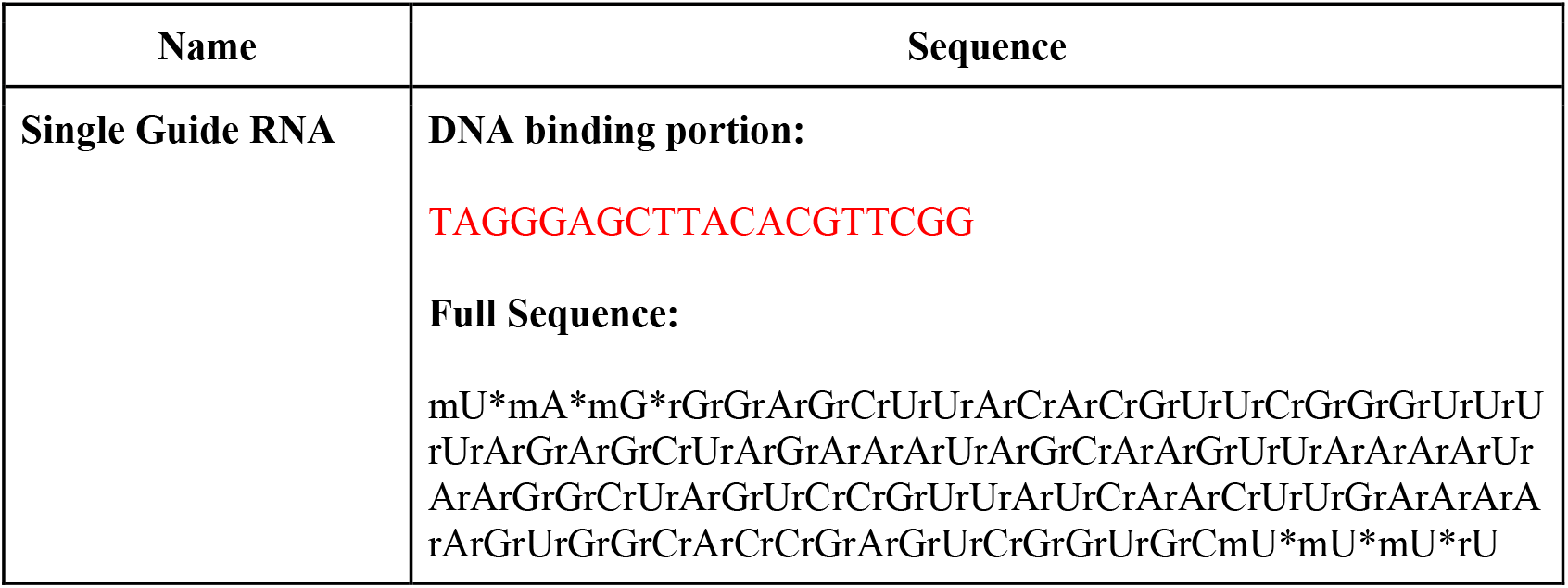

### 2.3 Method1 PROTOCOL

#### RT-PCR

First, hybridoma cells were homogenized using QIAshredder (Qiagen, 79654), and whole RNA was extracted using the Qiagen RNeasy Plus mini kit (Qiagen, 74134) by following the manufacturer’s protocol. Next, the sample was diluted to 100ng/μl in nuclease-free water and the RT-PCR reaction was performed as described by Meyer et al. [7]. 8μl of 100ng/μl RNA was added to 4μl dNTP (NEB, N0447S) and 4μl of 10 mM Reverse primer (mix 1). The samples were incubated at 72°C for 3 minutes. Next, the following solution was prepared (mix 2): 1μL of 80U/μL RNAse inhibitor (Invitrogen, 10777019), 2μL 100U/μL SMARTScribe reverse transcriptase (TAKARA, 639537), 7.8μL H2O, 8μL of 5x SMARTScribe buffer, 4μL of 20 mM DTT (TAKARA 639537), and 1.2μL of 100μM template-switch oligo [7]. 24μL of mix 2 was added to each tube of mix 1. In the thermocycler, the combined mix was incubated at 42°C for 60 minutes, then at 70°C for 5 minutes to stop the reaction. The reactions were held at 4°C. Lastly, the cDNA was purified using Select-a-Size DNA Clean & Concentrator Kit (ZYMO Research, D4080).

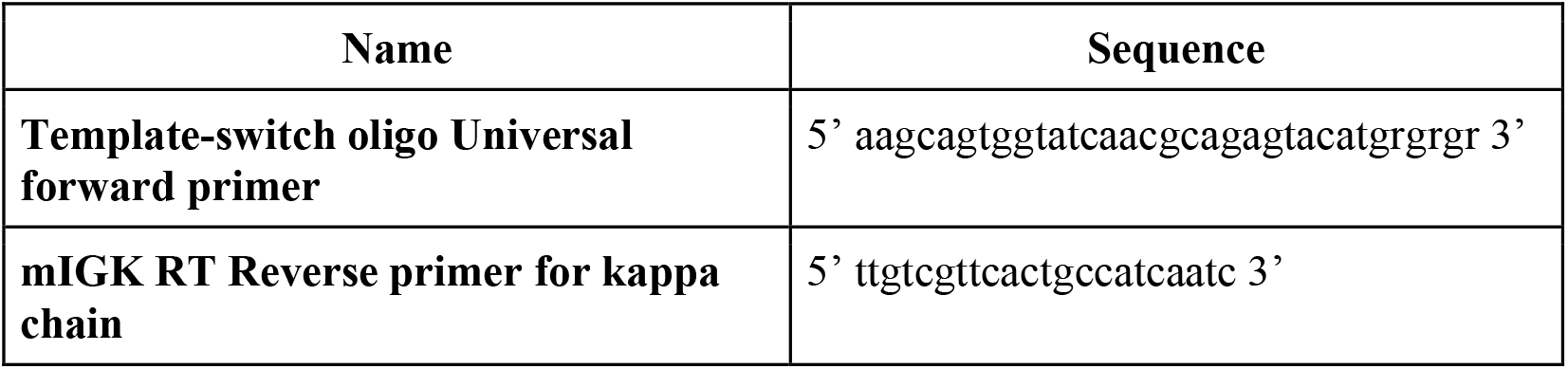

#### CRISPR Digestion

CRISPR/Cas9 digestion reaction was performed based on the following protocol: 3μl of the 10X CRISPR in vitro digestion buffer (20mM HEPES, 100mM NaCl, 5mM MgCl2, 0.1mM EDTA, pH 6.5), 1μl of 2μM sgRNA, 2μl of 1μM Cas9 protein, (Origene, TP790148), and 15ng of the cDNA (a molar ratio of 40:40:1) were added to a PCR tube and the volume was brought up to 30μl using nuclease free water. The reaction mixture was incubated at 37°C for 60 minutes followed by 65°C for 10 minutes to stop the reaction. The reaction was held at 4°C until the next step.

#### PCR

The PCR reaction was set up based on the following protocol: CRISPR treated cDNA, 2μl; Polymerase buffer 10X, 5μl; MgSO4, 2μl; dNTPs, 1μl; Reverse chain specific primer 10μM, 2μl; ISPCR Universal forward primer [7] 10μM, 2μl; Platinum™ Taq DNA Polymerase High Fidelity (Thermo Fisher, 11304029), 0.5μl; the total reaction volume was brought up to 50μl using water. The PCR reaction was performed based on the following settings, 94°C 90”, (94°C for 30’’, 56°C for 30’’, 72°C for 60’’) x30 cycles, 72°C for 7 minutes, the reaction was held at 4°C. PCR products were run on a 2% agarose gel at 100V for 30 minutes. 10μl of 100 bp Plus DNA Ladder (Biolab, N3200S) was used and bands at ∼560bp were cut and purified using the Qiagen PCR gel extraction kit (Qiagen, 28704).

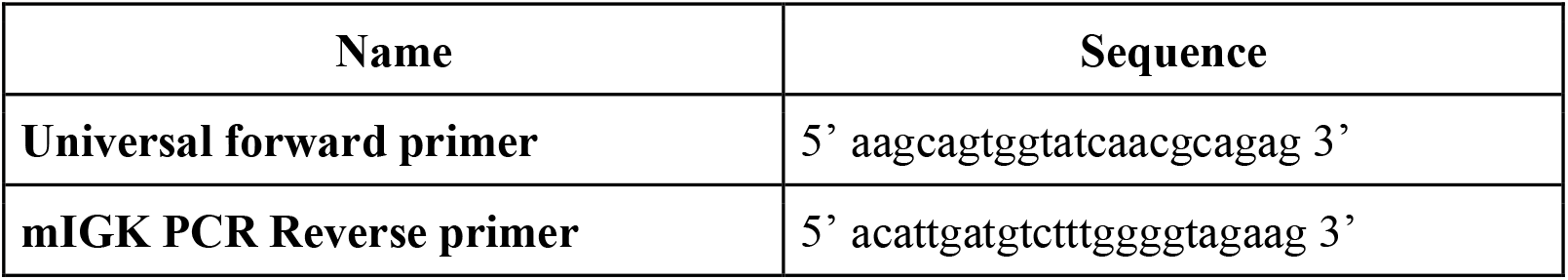

#### Cloning & Sequencing

The purified PCR product was then cloned using TOPO TA cloning kit (Thermo Fisher, 450641) according to the following protocol: PCR Product, 7μl; Salt solution (1.2M NaCl, 0.06M MgCl2), 1μl; TOPO vector, 1μl. The ligation was run for 30 minutes at room temperature. 5μl of ligation solution was used for the transformation. The transformation was done by heat shock. First, the tubes containing ligation and competent cells (Thermofisher, K4520-01), were incubated on ice for 30 minutes followed by incubation at 42ºC for 30 seconds immediately followed by incubation on ice for 2 minutes. Next, 250μl of SOC medium was added to the tube and incubated at 37ºC with shaking for 30 minutes. 100μl of the total transformation was plated onto LB agar plates containing carbenicillin (TEKNOVA, L1010) and 40μl of 40mg/ml X-gal solution (Invitrogen 15520-018F). The plates were incubated overnight at 37ºC. The next day, 12-16 white colonies were picked and incubated for 18 hours in 3ml LB broth-Carbenicillin at 37ºC. DNA extraction was performed the following day using the Qiagen miniprep kit (Qiagen, 27104). Samples were sent for Sanger sequencing (Ruifeng Biotech, Hayward, CA).

### 2.4 Method2 PROTOCOL

#### RT-PCR

RNA extraction and RT-PCR were performed based on the procedure described in Method1.

#### PCR

For the PCR reaction, 1ng of the purified cDNA was added to the PCR mixture. The PCR reaction was performed as described in Method1. The resulting product was purified using the Select-a-Size DNA Clean & Concentrator Kit.

#### CRISPR Digestion

The CRISPR/Cas9 digestion reaction was performed based on the following protocol: 3μl of the 10X CRISPR *in vitro* digestion buffer, 3μl of 2μM sgRNA, 6μl of 1μM Cas9 protein, 80ng (20:20:1 molar ratio) of the cDNA were added to a PCR tube, and the volume was brought up to 30μl using nuclease-free water. The reaction mixture was incubated at 37ºC for 60 minutes followed by 65ºC for 10 minutes. The reaction was held at 4ºC until the next step. 25μl of the reaction was run on a 2% agarose gel at 100V for about 30 minutes. 10μl of 100 bp Plus DNA Ladder (Biolab, N3200S) was used and bands at ∼560bp were cut and purified using the Qiagen PCR gel extraction kit (Qiagen, 28704).

#### Second PCR

The second PCR reaction was performed based on the procedure described earlier.

#### Second CRISPR Digestion

The second CRISPR/Cas9 digestion was performed like the first digestion.

#### Cloning & Sequencing

The purified ∼560 bp band was concentrated using Select-a-Size DNA Clean & Concentrator Kit. Cloning and sequencing were performed according to the procedure described in Method1.

### 3.0 RESULTS

#### 3.1 Proof of Concept of Specific cleavage of the SP2/0 Aberrant kappa Chain

In order to demonstrate the effectiveness and specificity of our sgRNA design. We cloned a functional kappa chain and the aberrant SP2/0 kappa chain into the pCR™ 2.1-TOPO™ TA vector using TOPO TA cloning kit (Thermo Fisher, 450641). We used the procedure described by Meyer et al [7]. Next, we transformed the vector into bacteria and grew the colonies in 3ml LB (as described in method1). Next, we performed DNA extraction and purification. The vectors were sent for sequencing to verify the sequences. IgBLAST and Nucleotide BLAST tools were used to identify the SP2/0 aberrant kappa chain (Sequence ID: FN422002.1) and the functional kappa chain. After sequence verification, we linearized the vectors using ScaI restriction enzyme (NEB, R3156S) (Fig. 3A). After gel purification, the samples were treated with CRISPR/Cas9. 3nM DNA template was used for each reaction. The sgRNA: Cas9: DNA molar ratios were 20:20:1 and the reactions were set up as described in Method1. 25μl of the reaction was run on a 1% agarose gel. The results are shown in Figure 3.

**Figure 2:**
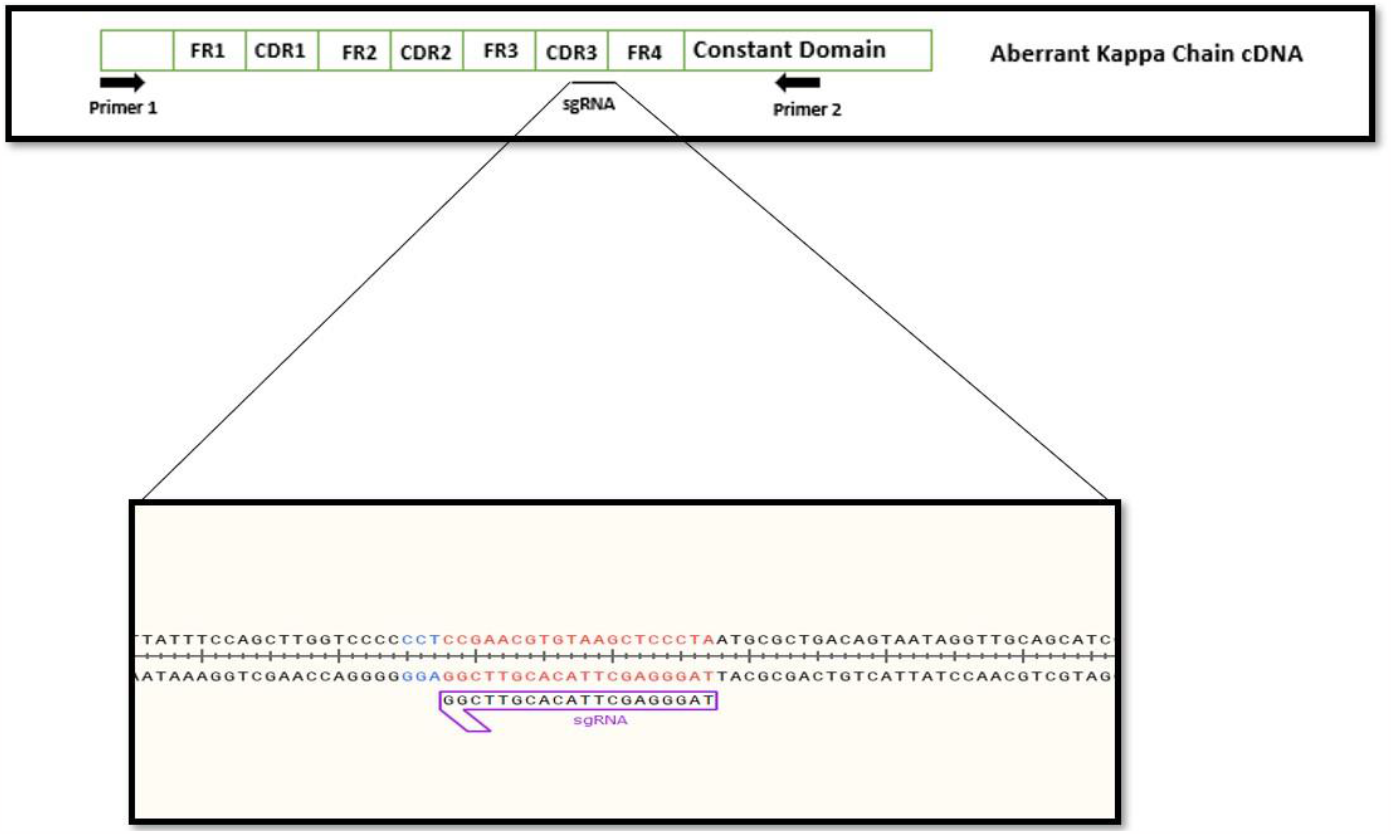
A schematic of the SP2/0 endogenous aberrant kappa chain cDNA including the primer binding sites and the sgRNA binding site (top image). Abbreviations: FR (frame work), CDR (complementary Determining Region), sgRNA (single guide RNA). The bottom image shows the sgRNA bindingitse, its sequence, and the PAM sequence (blue).

**Figure 3:**
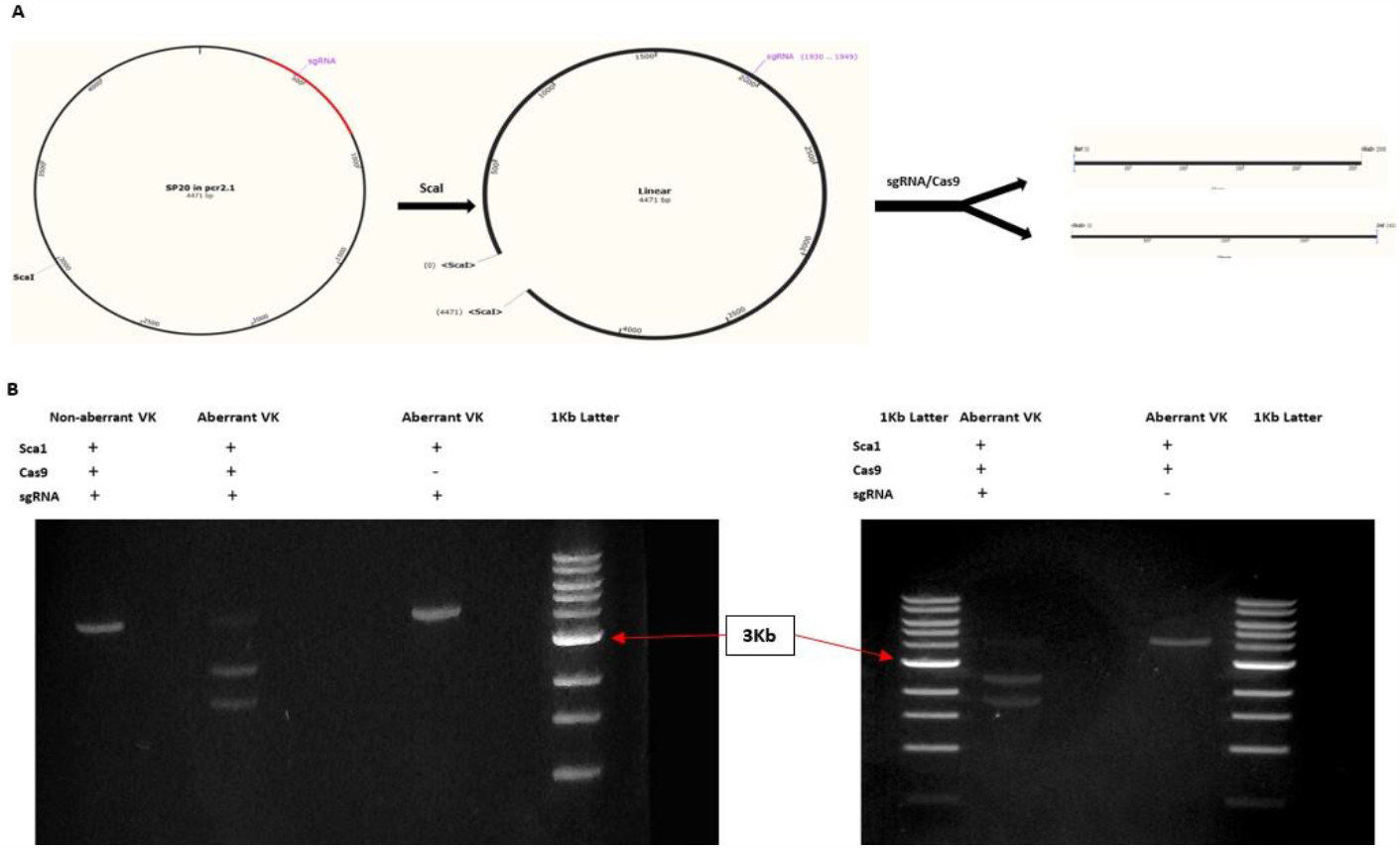
(A) shows the schematics of the proof of concept experiment. 4ug of the vector was first linearized using Seal (NEB) and rCutSmart buffer (NEB), the reaction is incubated for 1 hour at 37°C. The reaction then was run on a 1% agarose gel and purified. Next, the vector was digested using CRISPR/Cas9 (3μl 10X Cas9 buffer, l00dng DNA, 1 μl of 2μM sgRNA, 2μl of lμM Cas9, nucleases free water to bring up the total reaction volume to 30μl) the reaction was incubated at 37°C for 60 min followed by 10 minutes at 65°C. (B) shows the reaction products and appropriate controls run on a 1% agarose gel at 100V for 30 minutes.

This experiment demonstrated the sgRNA/Cas9 complex was able to specifically cleave the endogenous aberrant chain; thus, preventing its amplification during PCR. The result indicates that the aberrant chain vector was cleaved resulting in two smaller bands (approximately 2.5 Kb and 1.9 Kb) (Fig. 3B). The non-aberrant kappa chain bands remained a single non-cleaved band at approximately 4.4 Kb (Fig. 3B). When the aberrant chain vector was treated without Cas9 or sgRNA, there was no cleavage in the gel electrophoresis result, which demonstrates that both Cas9 and sgRNA were required for the specific cleavage at the CDR3 region. Our results also demonstrated that the designed sgRNA specifically targeted the aberrant chain without any significant off-target effects (Fig. 3B).

### 3.2 Method 1 (CRISPR Treatment After RT-PCR)

In this method, we treated the sample after performing the RT-PCR. As explained in the methods section, the molar ratio of sgRNA to Cas9 to the DNA template was 40:40:1. Then we PCR amplified the CRISPR/Cas9 treated cDNA. We cloned and sequenced two hybridoma clones using this method. The results showed a 24% to 25% decrease in the number of the aberrant chains in the treated versus non-treated samples (Fig. 4). IgBLAST and IMGT/V-QUEST [17] were used for sequence verification.

**Figure 4:**
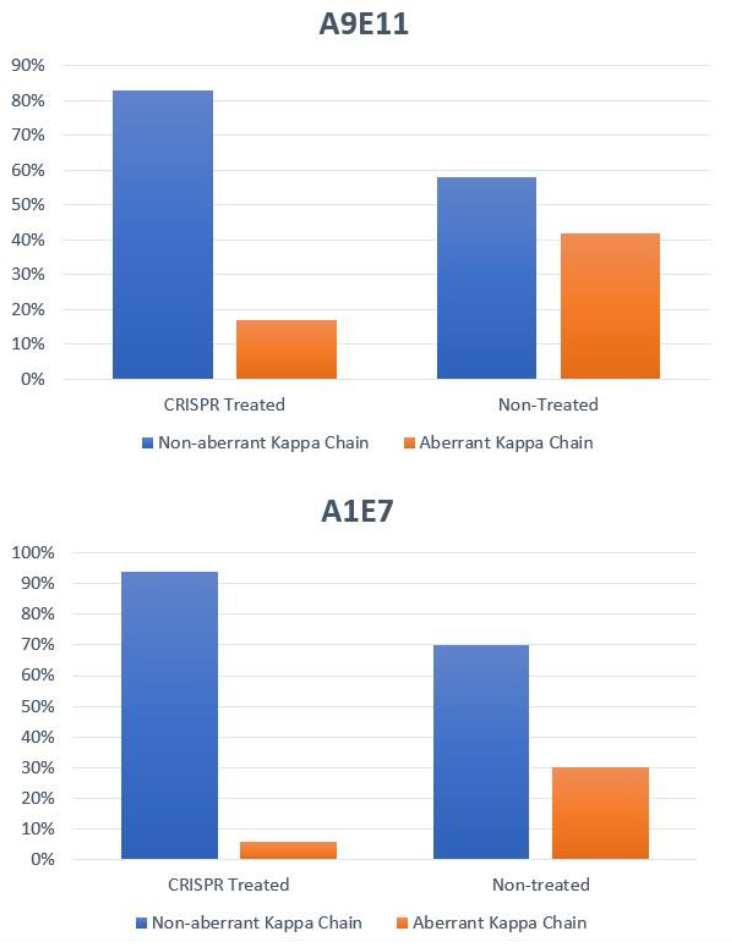
Shows the percentage of the aberrant versus non-aberrant kappa chain sequences in the CRISPR treated versus the non treated sample for clone A9E11 (top) and for clone A1E7 (bottom).

### 3.3 Method 2 (CRISPR Treatment of the PCR Amplicon)

In this method, we treated the PCR product with CRISPR/Cas9 with a 20:20:1 molar ratio as described in detail in the methods section. Next, we performed gel electrophoresis of the PCR product on a 2% agarose gel (Fig. 5A) and purified the 560 bp band. Next, we performed a second PCR reaction and treated the product with CRISPR/Cas9 again (double cut). It is important to note that the uncut band in gel 2 (Fig 5B) is much brighter than in gel 1 (Fig. 5A). This is because we have enriched the non-aberrant kappa chain transcripts. Next, we gel purified and cloned the 560 bp band shown in Figure 5B. The sequencing results showed an 88% decrease in the number of aberrant chain transcripts (Fig. 5C). IgBLAST and IMGT/V-QUEST [18] were used for sequence verification.

**Figure 5:**
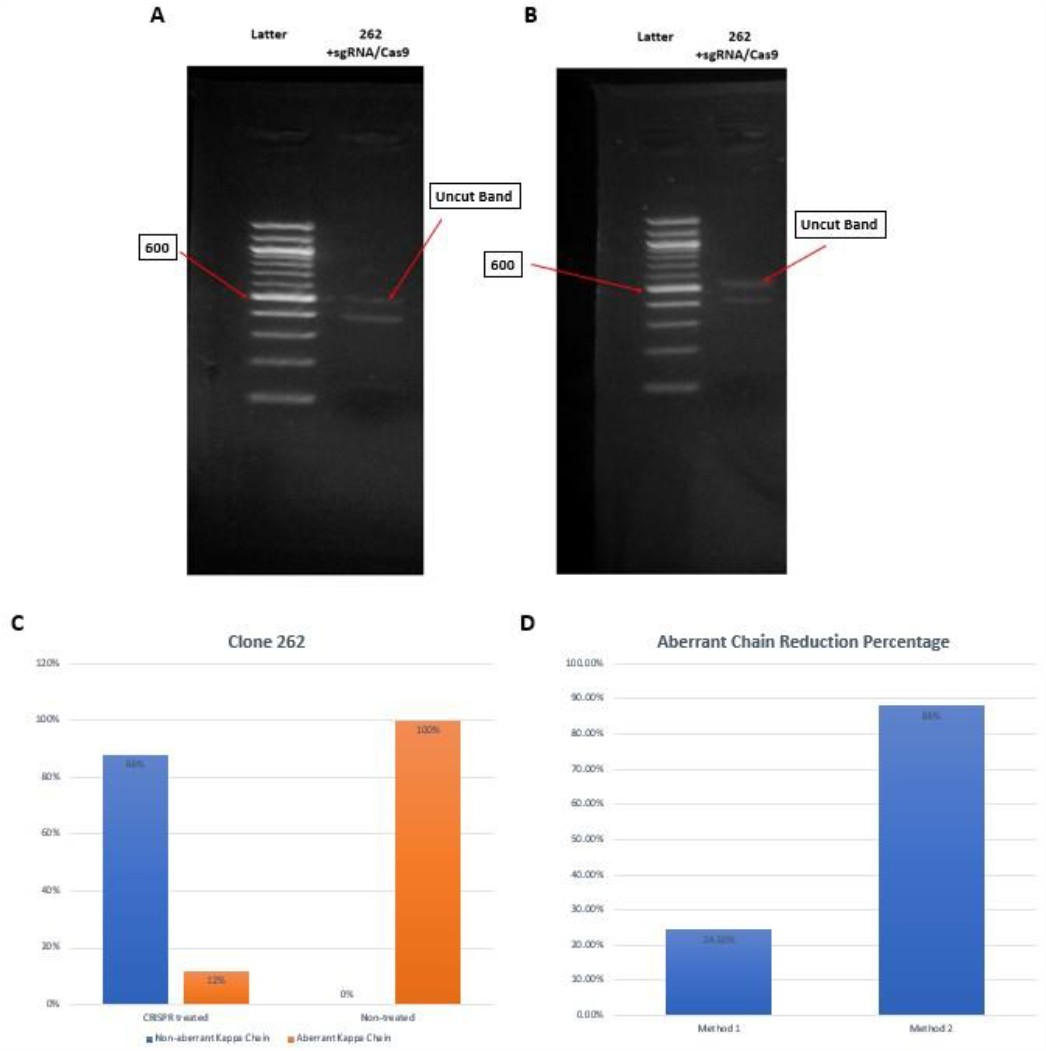
(A) shows the image of the CRISPR/Cas9 digested PCR product after the first digestion, the 560 bp band was purified and used as template for the second PCR. (B) shows the second PCR amplicon after the second CRISPR/Cas9 digestion. The 560 bp band was cloned and sequenced. (C) shows the sequencing results of the cloned fragment. This method decreased the percentage of the aberrant kappa chain transcripts by 88% in clone 262. (D) shows the comparison between method 1 versus method 2 in the percent reduction of the SP2/0 aberrant kappa chain sequences.

## 4.0 Discussion

Here, we demonstrated the effectiveness of using CRISPR/Cas9 to significantly reduce the number of aberrant kappa chain PCR amplicons that can interfere with sequencing analysis of hybridoma clones. The CRISPR/Cas9 digestion step took approximately 3 hours and the cost of the sgRNA, Cas9, and the digestion buffer was approximately $4 per sample. Additionally, compared to methods reported previously, our method provided multiple advantages. First, compared to the method reported by Juste et al [7], which uses restriction enzymes to cut the aberrant transcript, our method ensures a more specific cleavage of the aberrant kappa chain because it is possible that the restriction enzymes also cut the functional chain due to the unpredictable nature of the CDR3 region. Furthermore, we tried the method described by Yuan et al [9] for hybridoma clone 262. We did not observe any reduction in the number of the aberrant kappa chain transcripts (data not shown). In addition, we performed some optimization which increased this method’s efficacy from 24% (method 1) to 88% (method 2). Furthermore, we tried some commercially available Cas9 reaction buffers (NEB3.1 (NEB, B7203) and Origene Cas9 *in vitro* digestion buffer (Origene GE100053). We did not observe any significant difference in the cleavage efficiency. As previously described, in Method1, the sgRNA: Cas9: DNA molar ratio of 40:40:1 was used. For method2, we used a 20:20:1 molar ratio to keep the costs low. However, much higher molar ratios have been reported. For example, Prezza et al. reported a 35000:3500:1 to be an optimal ratio [13]. Furthermore, we only used a Cas9: sgRNA ratio of 1:1 as optimal because it has been reported frequently in various CRISPR/Cas9 based *in vitro* digestion protocols (New England Biolabs and Origene). Changing the sgRNA: Cas9 ratio without changing the DNA concentration may also help improve the efficiency of our method. In addition, further optimization of our method could include testing other Cas nucleases [19]. For example, high-fidelity Cas versions with increased fidelity and thermostability [14,20] could also increase the aberrant chain depletion efficiencies.

## 5.0 CONCLUSION

In conclusion, in this paper we report the development of a cost effective method to cleave the SP2/0 endogenous kappa transcript using the CRISPR/Cas9 system. In order to ensure the specific cleavage of the SP2/0 aberrant chain, we designed our sgRNA to target the CDR3 region of the transcript. Lastly, the experiments demonstrated the effectiveness of using CRISPR/Cas9 to significantly reduce the number of the SP2/0 derived endogenous aberrant kappa chain sequences especially after optimization of method1.

## 6.0 ACKNOWLEDGEMENTS

The experiments were designed by Armin Ahnoud and Chong He. The experiments were performed by Armin Ahnoud. The manuscript was written by Armin Ahnoud and Chong He. Chong He, Michael Strainic, and Wenwu Zhai edited the manuscript.

